# Rapid Regulation of Local Temperature and TRPV1 Ion Channels with Wide-Field Plasmonic Thermal Microscopy

**DOI:** 10.1101/2022.06.28.497933

**Authors:** Rui Wang, Jiapei Jiang, Xinyu Zhou, Zijian Wan, Pengfei Zhang, Shaopeng Wang

## Abstract

Plasmonic absorption of light can create significant local heat and has become a promising tool for rapid temperature regulation in diverse fields, from biomedical technology to optoelectronics. Current plasmonic heating usually relies on specially designed nanomaterials randomly distributed in the space and hardly provides uniform temperature regulation in a wide field. Herein we report a rapid temperature regulation strategy on a plain gold-coated glass slip using the plasmonic scattering microscopy, which can be referred to as wide-field plasmonic thermal microscopy (W-PTM). We calibrated the W-PTM by monitoring the phase transition of the temperature-sensitive polymer solutions, showing that it can provide a temperature regulation range of 33-80 °C. Moreover, the W-PTM provides imaging capability, thus allowing the statistical analysis of the phase-transitioned polymeric nanoparticles. Finally, we demonstrated that W-PTM can be used for noninvasive and local regulation of the transient receptor potential vanilloid 1 (TRPV1) ion channels in the living cells, which can be monitored by simultaneous fluorescence imaging of calcium influx. With the nondestructive local temperature-regulating and concurrent fluorescence imaging capability, we anticipate that W-PTM can be a powerful tool to study cellular activities associated with cellular membrane temperature changes.

---

Temperature is a fundamental environmental parameter that is critical in cellular activity regulations. The classical temperature control techniques employ heating sources to conduct the heat transfer, which can be time-consuming and nonuniform over the target surface. In recent decades, plasmon resonances in nanostructures, especially metallic nanoparticles, have been proved to be efficient in regulating localized heat rapidly and feasibly^1,2^. These nanometer-sized heaters are capable of harvesting light due to the internal decay of hot carriers, facilitating many practical applications, such as photothermal therapy^3^, neuron activation^4^, phase separation^5,6^, gas sensing^7^, and heterogeneous catalysis^8,9^. However, localized heating achieved by metallic nanoparticles still suffers from poor space precision of heating due to the random distribution of metallic nanoparticles, resulting in bulk heating on the sample. Moreover, the nanocavities created by the tightly positioned metal nanoparticles may also generate excessive heat, leading to overheating on the target^10,11^.

Here we show a wide-field plasmonic thermal microscopy (W-PTM), which provides rapid temperature regulation and uniform temperature distribution over its detection field (Figure 1a). Specifically, W-PTM can utilize the evanescent properties of surface plasmonic waves to limit the heating space within ~100 nm nearby the gold surface, providing a feasible way to selectively heat the temperature-sensitive membrane proteins, such as the transient receptor potential vanilloid 1 (TRPV1) ion channels, for regulating the cell activities^12,13^. First, we calibrated the temperature regulation range and temporal dynamics of W-PTM by monitoring the phase transitions of temperature-responsive polymers in aqueous solutions^14,15^. Then, we employed W-PTM to selectively activate ion channels in TRPV1 transfected HEK-293T cells, accompanied by fluorescence imaging to monitor the intercellular calcium ions influx processes, demonstrating the feasibility of using W-PTM to study the temperature responsiveness of living cells.

**Figure 1.**
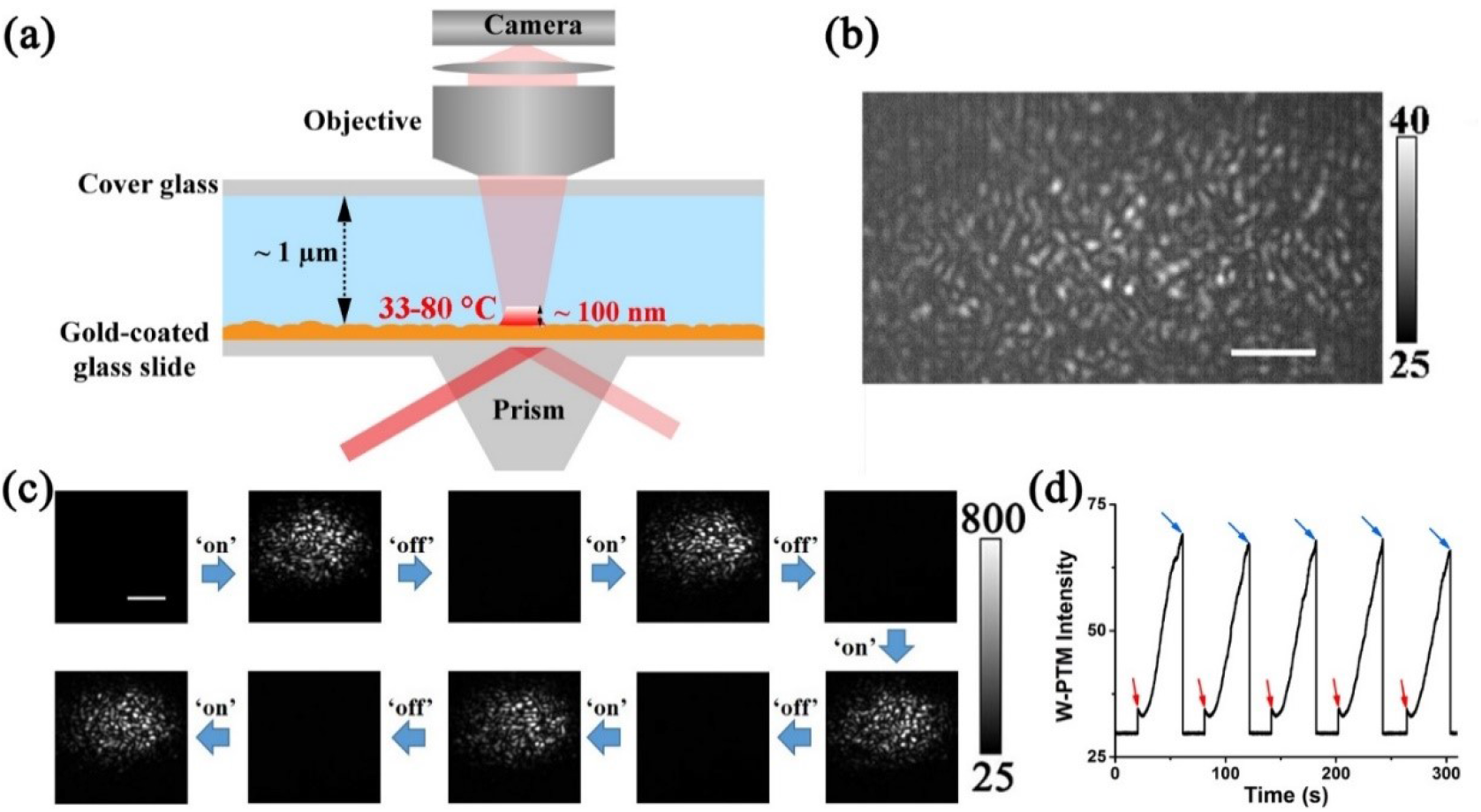
(a) Schematic overview of the experimental set-up of W-PTM. The thickness of the flow channel is ~100 μm. The p-polarized 660 nm laser was directed to the Au chip to reach SPR. (b) W-PTM image of roughness scattering from Au chip itself under 1.33 kW/cm^2^. (c) W-PTM images of excitation power-controlled multiple switching process of polymer phase transition. 1 mg/mL HPC (LCST = 54 °C) was flowed to the Au chip at a rate of 0.5 mL/mL. Excitation power densities for ‘on’ and ‘off’ states were 2 kW/cm^2^ and 1.33 kW/cm^2^, respectively. Scale bar, 5 μm. (d) Corresponding ensemble W-PTM intensity change in (c). The red and blue arrows indicate the switch of the laser power density between 2 kW/cm^2^ and 1.33 kW/cm^2^, respectively.

The W-PTM was developed based on the plasmonic scattering microscopy (PSM)^16–18^, a novel wide-field surface plasmon resonance (SPR) imaging approach. In addition to the wide-field imaging ability, which is beyond the capabilities of localized SPR detection with metallic nanoparticles, the W-PTM also shares the advantages of PSM over the traditional SPR systems: 1) the W-PTM does not record the propagating plasmonic waves with long decaying length along the surface into the images, thus providing high spatial resolution and Gaussian-distributed point spread function for automatic image processing with conventional open-source software such as ImageJ. As a result, W-PTM enables easy monitoring of thermal dynamics in the time domain by tracking the formation of polymeric aggregation (i.e., phase transition) particles. 2) W-PTM does not record the strong reflection, thus allowing the incident intensity up to 3 kW/cm^2^ so that a wide temperature regulation range can be achieved from room temperature to ~80 °C. 3) W-PTM records the light from the top of the gold surface, making it possible to integrate with fluorescence detection approaches, whose signals are massively dissipated through the gold surface in traditional SPR systems^19,20^.

To investigate the local heating behavior on W-PTM, polymers with low critical solution temperature (LCST) were chosen due to their phase transition and non-photobleaching features^14,15^. Moreover, a wide range of responsive temperatures (i.e., LCST) can be easily achieved by applying the salt effect to a specific polymer^21,22^. Temperature-dependent dynamic light scattering (DLS) results showed that 8 groups of LCST polymer precursor with LCST ranging from 33-80 °C could be achieved by dissolving hydroxypropyl cellulose (HPC) or poly(ethylene oxide)(PEO, Mw = 100, 000) in Na2HPO4 or NaCl aqueous solution with various salt concentrations (Figure S1). As shown in Figure 1, after applying a specific LCST polymer precursor, such as 1 mg/mL HPC (LCST = 54 °C), to the Au chip surface, dynamic formation of bright scattering spots can be observed in the W-PTM image when incident light power above a threshold level, in contrast with weak roughness scattering from Au chip itself (Figure 1b). We also confirmed that the appearance and disappearance of these bright spots can be switched ‘on’ and ‘off’ reproducibly for 5 times by tuning the excitation power of the W-PTM (Figure 1c, d, Figure S2). These phenomena suggest that these bright spots shown in the scattering images are from dynamic formation of polymer nanoparticles associated with local temperature change.

In order to further confirm the feasibility of LCST polymer in sensing the local temperature of W-PTM, we tested the phase transition processes of four LCST polymer precursors (LCST = 33, 45, 62, 72 °C, referred to as 33 °C LCST, 45 °C LCST, 62 °C LCST, 72 °C LCST, respectively) on three independent Au chips, respectively. By measuring ensemble image intensity as a function of excitation power density, we found that the phase transitions of these LCST polymers are consistent among these chips. Interestingly, HPC-based (LCST = 33, 45 °C, Figure 2a) and PEO-based (LCST = 62, 72 °C, Figure 2b) LCST polymers show inverse correlation between W-PTM intensity and LCST temperature. Specifically, W-PTM intensity produced by 33 °C LCST is higher than that produced by 45 °C LCST under the same power excitation, while the opposite trend is found on 62 °C LCST and 72 °C LCST. We attribute the former to a higher phase transition threshold energy required for 45 °C LCST than 33 °C LCST and thus resulting in a slower and smaller nanoparticle formation at the same power, which can also be reflected by the slower kinetics of the W-PTM intensity change of 45 °C LCST. However, the ensemble signal analysis cannot explain the difference between 62 °C LCST and 72 °C LCST, for which single-particle analysis is required.

Next, we calibrated the surface temperature of the gold chip using these LCST polymers. We gradually increased the W-PTM excitation power density in a step-by-step manner while monitoring the phase transition of a specific LCST polymer. As shown in Figure S3, the W-PTM intensity becomes very sensitive to the excitation power when the local temperature reaches the phase transition temperature, defined by the W-PTM intensity change exceeding five times the fluctuation of the background signal (Figure S3c, Video S1, S2). And its equilibrium temperature was defined as the phase transition temperature of the specific LCST polymer. Figure 2c shows the local equilibrium temperature under various power densities by measuring these LCST polymers. Typically, local heat would reach equilibrium within 10 s of excitation, reflected by the steady increase of the W-PTM intensity. We found that the local equilibrium temperature is exponentially related to the power density. Our results clearly demonstrate that a local temperature of ~62 °C can be reached at 1.5 kW/cm^2^, a power density often used in the single-molecule analysis. The localized heat of W-PTM may cause the inactivation of thermally unstable samples, such as proteins^23^, etc. Therefore, our temperature calibration curve helps to find out the up-limit of the power density that can be used to detect heat-sensitive samples.

**Figure 2.**
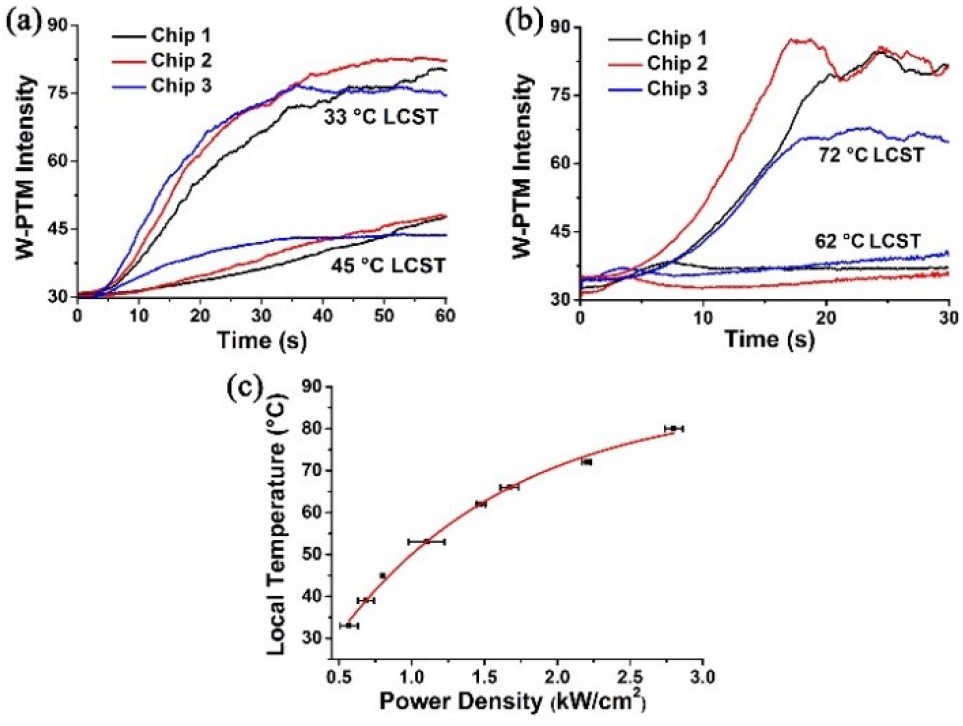
(a, b) Ensemble W-PTM intensity as a function of excitation time of four groups of LCST polymers (33 °C LCST, 45 °C LCST, 62 °C LCST, 72 °C LCST) on three Au chips independently. The excitation power density of (a) and (b) is 1.33 kW/cm^2^ and 3 kW/cm^2^, respectively. (c) Localized equilibrium temperature against power density calibrated by various LCST polymers. The fitted curve shows an exponential relationship between equilibrium temperature and excitation power density (r^2^ = 0.995).

To validate the above findings, low-density lipoprotein (LDL)^24^, a natural heat-sensitive protein with a transition temperature at ~ 78 °C, was used as a reference. We determined the phase transition power density and the corresponding phase transition time of LDL to be 2.1 kW/cm^2^ and 16 s, respectively, which is comparable to that of 80 °C LCST (2.5 kW/cm^2^ and 19 s). As shown in Figure S4, both LDL and 80 °C LCST showed phase transition behavior when excited at the same power density (3 kW/cm^2^), but the ensemble W-PTM intensity change of LDL is close to an exponential increase, while that of 80 °C LCST is closer to linear increase (Figure S4e). These dynamics can be explained by the slight lower phase transition temperature of LDL making burst nucleation of LDL at this power density, while the 80 °C LCST is just reaching the phase transition temperature and polymerizing at a slower rate. Nevertheless, this result confirms that local temperature calibration using LCST polymer is reliabile and accurate.

Taking advantage of the single-particle analysis capability of W-PTM, an image processing algorithm was employed to analyze the size change of the generated particles during the phase transition (details can be found in Supporting Information). We first subtract a previous frame from each frame to remove background features, resulting in the differential images (Figure 3a). Then these differential images were analyzed by Trackmate, a plugin software affiliated with ImageJ, to obtain the showing time, location and W-PTM intensity of each particle. Meanwhile, according to the intensity-size equation we built previously^16^ (details can be found in Supporting Information), the W-PTM intensities are correlated with their actual > size. By plotting cumulative particle numbers against size and time, we can obtain the phase transition dynamics of LCST polymers at the single-particle level. For example, nanoparticles with a size centered at ~150.9 nm will appear for 33 °C LCST. polymer after excitation at 1.33 kW/cm^2^ for about 20 s, and, while the number of particles increases, the particle size remains unchanged during the entire phase transition process (Figure 3b, c). In addition, we also found that 45 °C LCST (Figure 3d, e) yielded smaller particle sizes (~108.8 nm) than 33 °C LCST under the same conditions with similar counts, which is consistent with ensemble measurements on 33 °C LCST and 45 °C LCST (Figure 1d). In the case of 62 °C LCST and 72 °C LCST, single particle analysis indicated that the former has larger particle size (130.2 nm vs. 88.4 nm) (Figure 3f-i), but 72 °C LCST showed faster particle generation kinetics, which resulted in twice as many particles as 62 °C LCST. This phenomenon can explain why 72 °C LCST has a stronger ensemble W-PTM intensity than 62 °C LCST (Figure 2b). Overall, single particle analysis can provide more information on the phase transition process than ensemble signal analysis and provide phase transition details at the nanometer scale.

**Figure 3.**
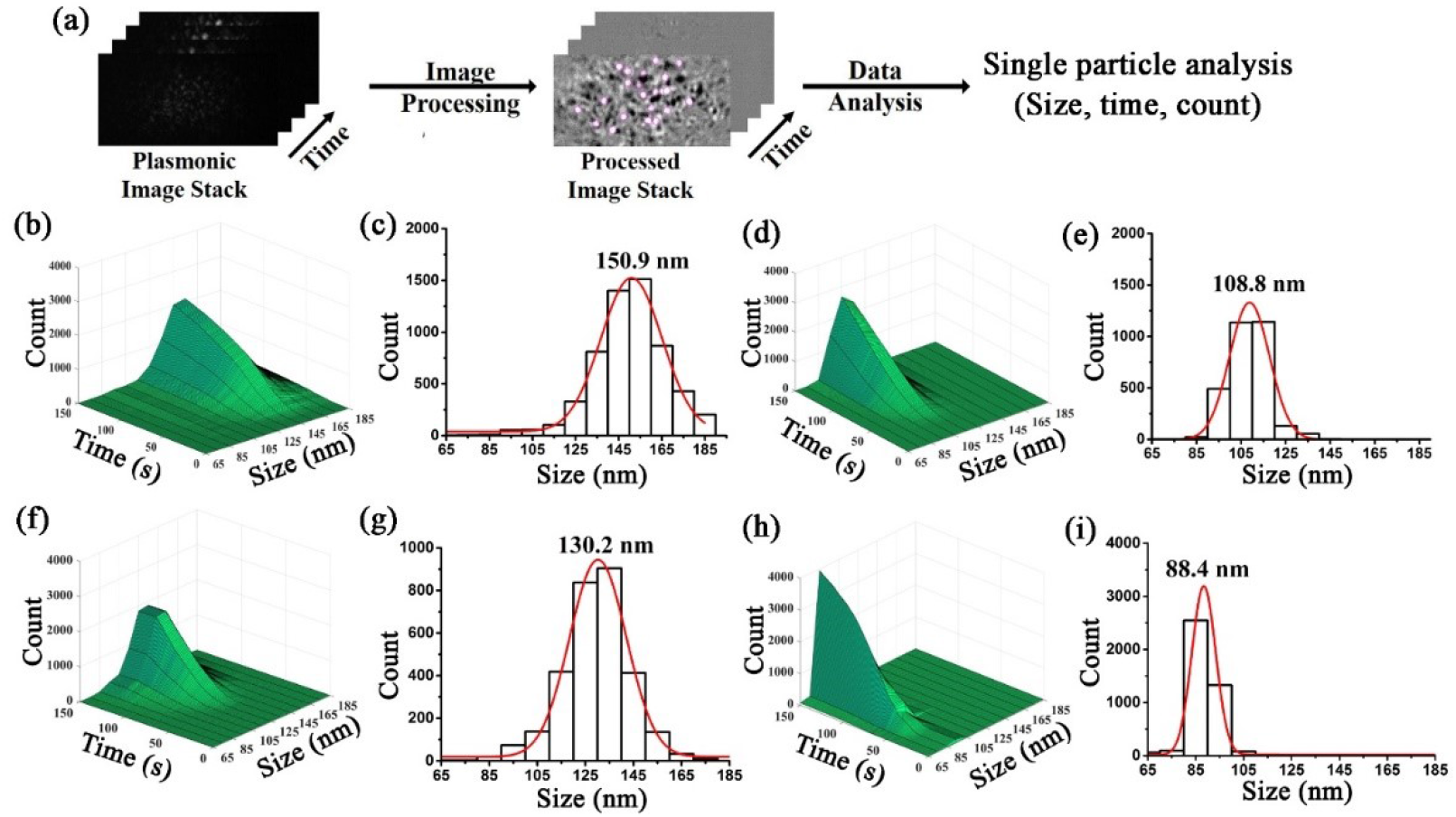
(a) Single-particle analysis processes of W-PTM data. A detailed description of the image processing and data analysis can be found in the Supporting Information. (b, d, f, h) Cumulative particles numbers versus size and time of 33 °C LCST (b), 45 °C LCST (d), 62 °C LCST (f) and 72 °C LCST (h). The excitation power density for (b, d) and (f, h) is 1.33 and 3 kW/cm^2^, respectively. (c, e, g, i) The size histograms of (b, d, f, h) at 100 s, respectively. The solid lines are Gaussian fittings for phase transition generated nanoparticles.

We expect W-PTM to be applied to biological micro-heaters, owing to the advantages of non-invasiveness, label-free feature, accurate temperature control, and light-triggered fast switching feature. As a proof of concept, we attempted to exploit the local thermal effect of W-PTM to selectively activate transient receptor potential vanilloid 1 (TRPV1) ion channels. The TRPV1 ion channel is a calcium-permeable non-selective cation channel that can be activated by capsaicin, heat (> 42 °C), pH (< 5.9), voltage, and other stimuli. After activated, the action potential can be triggered to promote downstream intracellular signal transduction processes^12,13^. Thus elucidating how TRPV1 responds to heat or drugs is vital to understanding diseases that affect every major organ system of the body. However, the activation of TRPV1 is often limited to a few ensemble triggering methods, such as halogen lamps, focused lasers, and drug flows, making it difficult to achieve selective activation^25,26^.

To investigate the response of TRPV1 ion channels to the localized heat of the W-PTM, we employed the calcium imaging by integrating a set of fluorescence (FL) imaging pathways along with W-PTM (Figure 4a, Figure S5), and the TRPV1-transfected cells were also stained by Fluo-4AM^27^, a calcium indicator dye that responds to TRPV1 activation by changes in FL intensity (Figure S6, S7). From the superposition of the FL image and the W-PTM image (Figure 4b), we can see that only a part of the TRPV1-transfected cells can be thermally stimulated. Next, we monitored the FL changes of these cells responding to W-PTM excitations. As can be seen in Figure 4c and Video S3, we found that when the equilibrium temperature (33 °C) provided by the W-PTM (power density = 0.5 kW/cm^2^) was lower than the activation threshold temperature of TRPV1, both heated and non-heated cells remained un-activated, characterized by a continuous decrease in fluorescence (Figure 4d), corresponding to the photobleaching of the Fluo-4AM. Then, we further increased the power density of W-PTM to 1.2 kW/cm^2^, with an equilibrium temperature ~12 °C higher than the activation threshold of TRPV1. We found that the FL intensity of heated cells showed a sudden drop but increased gradually. We suspect that the sudden drop in FL in heated cells is related to quick photobleaching of Fluo-4AM due to the onset of W-PTM excitation, while the subsequent FL increase can be explained by the large influx of calcium ions accompanying the opening of the TRPV1 ion channel. Meanwhile, the unheated cells remain un-activated, characterized by consistent photobleaching (Figure 4e, f, Video S4). These results show that the localized thermal effect of W-PTM can achieve precise and controllable activation of TRPV1 ion channels, which could be use for functional study of TRPV1 and accelerate related drug discovery process.

**Figure 4.**
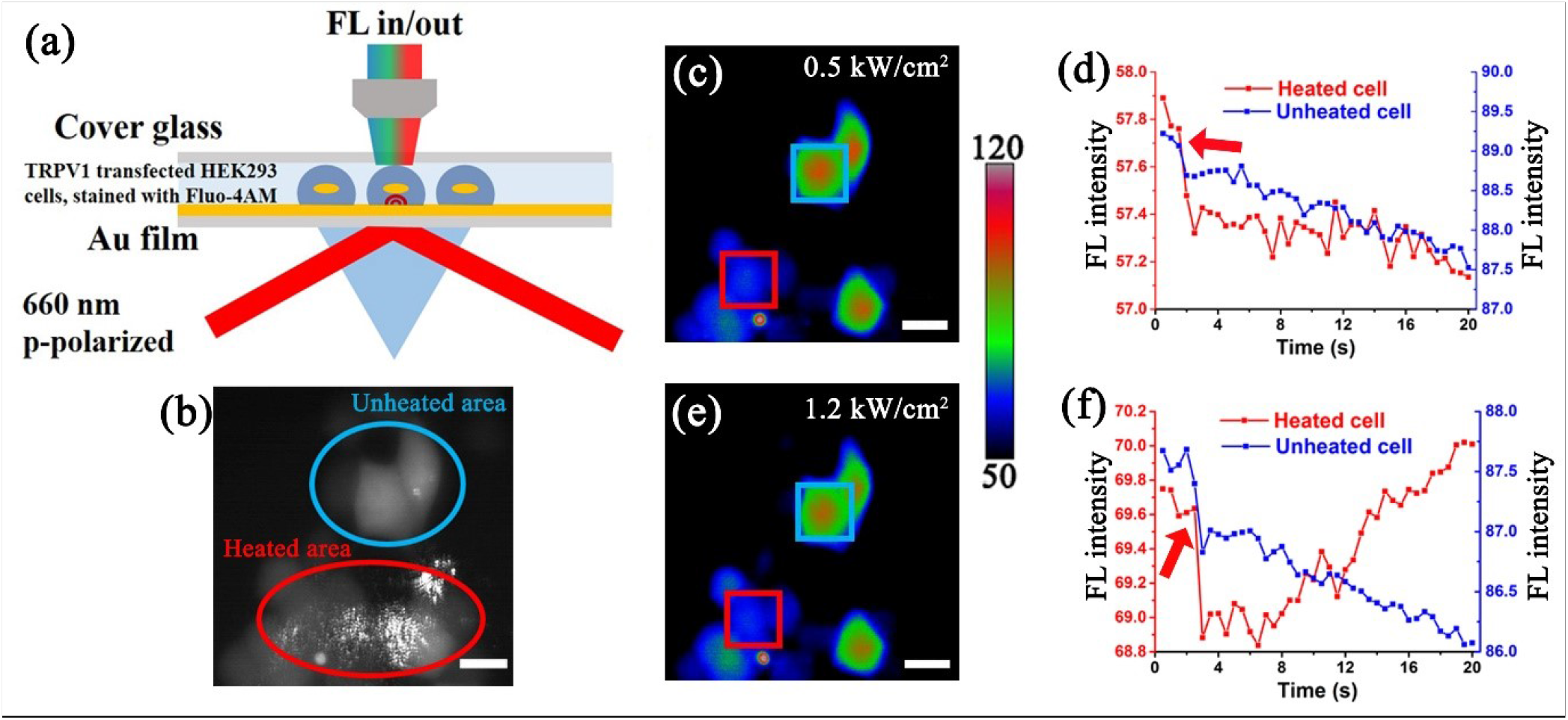
(a) A sketch of imaging set-up for selective TRPV1 activation monitoring. A more detailed set-up can be seen in Figure S4. (b) An overlap of the FL image and the W-PTM image, showing the locations of activated cells (red circle) and unactivated cells (blue circle). (c, e) The FL imaging snapshots (pseudo-color) of cells excited at 0.5 kW/cm^2^ (c) and 1.2 kW/cm^2^ (e). Cells of interest in the W-PTM view and out of W-PTM view were marked with the red and blue square, respectively. (d, f) the corresponding FL intensity traces of the cells of interest in (c, e), respectively. The red arrow marked the time when W-PTM excitation started.

Compared with the traditional micro-heating methods, such as plasmonic metallic nanoparticles heating, W-PTM provides a more precise and controllable micro-region heating. Owing to the evanescent properties of surface plasmonic waves on W-PTM, the Z-dimension of heating space is limited to ~100 nm nearby the gold surface, while the XY-dimension can be controlled by the laser focus ranging from ~1 to hundreds μm^2^. Moreover, the local temperature on the W-PTM is controllable within 33-80 °C with no overheating effects. Meanwhile, W-PTM can also analyze thermal transition kinetic processes at the single-molecule or single-particle scale, such as the phase transition process of LCST polymers. In addition to quantify the particle size of these generated nanoparticles, W-PTM can also provide 1 ms temporal resolution for rapid real time counting of the particles, providing more information than ensemble measuring methods like DLS.

Until now, the heat activation of TRPV1 is often limited to ensemble triggering methods that lack of selectivity at cell level. In this work, we applied W-PTM in selective heating of the TRPV1 transfected cells, while calcium fluorescence imaging was used as a reference. We demonstrated that W-PTM can selectively activate the cells in the field of view (as few as several cells) in an excitation power-dependent manner without affecting surrounding cells. Our previous study demonstrated that PSM, the parent technique of W-PTM, can image cell deformation at the single focal adhesion level. Therefore, in future studies we could monitor the dynamic cell deformations to characterize the thermally controlled ion channel states. We anticipate that W-PTM will become a powerful tool for studying TRPV1 and other thermal responsive cellular activities, and accelerating drug screening throughput in the future.

## ASSOCIATED CONTENT

### Supporting Information

Experimental methods, Temperature-dependent DLS results of LCST polymers, W-PTM images of LCST polymers and LDL, Optical design of W-PTM set-up coupled with FL imaging, Transfection efficiency optimization, FL images of TRPV1 transfected cells triggered by capsaicin, Single-particle analysis code (PDF)

Video S1 (avi)

Video S2 (avi)

Video S3 (avi)

Video S4 (avi)

## AUTHOR INFORMATION

**Corresponding Author**

*Pengfei Zhang, *Shaopeng Wang

## Author Contributions

S. W., P. Z. and R. W. conceived the project; R. W. and J. J. performed the experiment; J. J. conducted the single-particle analysis; X. Z. helped with cell experiments including cell culturing and plasmid transfection; Z. W. fabricated the Au chips; P. Z. built the W-PTM setup and S. W. acquired funding. The manuscript was written through contributions of all authors. All authors have given approval to the final version of the manuscript. §These authors contributed equally.

## Notes

The authors declare no competing financial interests.

## ACKNOWLEDGMENT

We are grateful for financial support from the National Institute of General Medical Sciences of the National Institutes of Health grant R01GM107165. We acknowledge the use of facilities within the ASU NanoFab supported in part by NSF program NNCI-ECCS-1542160.

## Notes

### Competing Interest Statement

The authors have declared no competing interest.

