## Supplemental Figure 1-7 for "Rapid Regulation of Local Temperature and TRPV1 Ion Channels with Wide-Field Plasmonic Thermal Microscopy"

**Methods**

**LCST polymer precursor solution preparation**

Hydroxypropyl cellulose (HPC) and Poly (ethylene oxide) (PEO) stock solution at a concentration of 10 mg/mL and 2.36 mg/mL was prepared, respectively. Then a group of LCST polymer precursor solutions were prepared by mixing the stock solution with proper concentration of $Na_2HPO_4$ or NaCl. The detailed information can be seen in Figure S1.

**W-PTM monitoring phase transition of LCST polymer**

LCST polymer precursor solutions are fed into the system in the format of gravity pump system. Solutions are allowed to flow for a few seconds before the laser light was turned on to balance out the pressure difference. As soon as the excitation laser was turned on, W-PTM recording starts simultaneously. Local heat from focused laser built up and reach equilibrium with heat dissipate, reaching a final stabilized temperature, which cause the phase transition of the polymer solution when surpassing the LCST threshold temperature. The precipitated aggregated polymeric particles that diffuse and land onto the gold chip are recorded as bright spots in the W-PTM images. The exposure time was power density-dependent as we had to avoid the overexposure of the camera. Typically, an exposure time of 0.1 ms was used when the power density was below 1 $kW/cm^2$; while the exposure time was shorten to 0.03 ms with a neutral density (ND) filter (ND 0.6) applied when the power density was between 1 and 2 $kW/cm^2$; and the exposure time of 0.1 ms and ND 2 filter was used for power density above 2 $kW/cm^2$.

To find out the equilibrium temperature at a specific W-PTM power density, we monitored the W-PTM intensity change of these LCST polymers while gradually increasing the excitation power density in a step-by-step manner. Typically, we started with 0.33 kW/cm$^2$, and an increment of 0.33 kW/cm$^2$ was conducted if the W-PTM intensity change did not exceed five times the background fluctuation in 30 s. In particular, to achieve a higher temperature sensing precision, a smaller step size (like 0.1 kW/cm$^2$) was applied when it came close to the equilibrium temperature.

**TRPV1 ion channels transfection**

HEK 293T cells (ATCC, Product Number: CRL-11268) were transfected with TRPV1 and dTomato gene-containing plasmid (addgene, Product Number: #140945) with the help of Lipofectamine™ 3000 Transfection Reagent (Invitrogen™, Product Number: L3000075). All transfection procedures followed the manufacturer's protocols. After repeated transfection twice, the total transfection efficiency was estimated to be 80-90% (Figure S6). To prove the functionality of the TRPV1 ion channels, we performed capsaicin triggering, which is a gold standard for testing TRPV1. First, we stained these cells with Fluo-4AM: Fluo-4AM (Abcam, CAS Number: 273221-67-3) stock solution (1mM in DMSO) was prepared and diluted in HBSS at a concentration of 10 μM. Then 500 μL of the above Fluo-4AM HBSS solution was gently added to the incubation well of the cells (Volume ratio of dye and cells is 1:4), and incubated with cells for 50 min at room temperature, then kept in an incubator for 30 min at 37 °C. Then, these stained cells were washed with HBSS to remove the excess dye and placed on a microscope for fluorescence monitoring. After capsaicin (Sigma-Aldrich, Product Number: M2028) to a final concentration of 30 μM, an apparent fluorescence enhancement can be observed under 488 nm excitation, indicating the activation of TRPV1 (Figure S7).

The bright-field and fluorescence images were taken on an inverted microscope (Olympus IX81) equipped with a halogen lamp and a 40X objective (NA 0.75). The 525/10 nm band-pass filter was used in the excitation route to excite dTomato, a fluorescence indicator incorporated in the TRPV1 plasmid. And 488/10 nm band-pass filter was used to excite Fluo-4AM.

**Single-particle analysis**

For precipitated single particles that landed on the gold chip, a light spot will appear in the processed image. Such bright spot can be captured and analyzed by the Trackmate tool in ImageJ. Threshold are set that the averaged particle number per frame is ~1. Using home built Matlab based algorithm, the intensity, location, appearing time information will be categorized and analyzed. Size of the precipitated particle is converted by correlating the intensity of the light spot to the size-intensity calibration curve. For temporal dynamic analysis of the precipitated particles, the particles are clustered into bins of every 1 second and 5 nanometers. And the Size-Time-Count multi-dimensional curve is obtained based on the resulting countings.

The diameters of the aggregated polymeric nanoparticles were determined using the calibration curve established previously (Fig. 2g, Nature methods, 2020, 17(10): 1010-1017). Given the fact that the calibration curve was based on polystyrene, which shares similar refractive index (1.58) with HPC (1.54, https://www.guidechem.com/trade/hydroxypropyl-cellulose-id382703.html) and PEO (1.46, https://www.guidechem.com/encyclopedia/poly-ethylene-glycol--dic21252.html), no further cross-section adjustment was conducted.

**Matlab code of single-particle analysis**

```
clear all;
close all;

directory_name = 'G:\20210524 Chip 3\New folder\1\';
filename = 'Result';
fps = 69;
power = 45;
exposure_time = 0.2;
factor = 37.5*2*100/(power*exposure_time);
timeframe = fps * 70;
sectionnumber = timeframe/fps;



for f = 1:1:1
    clear data01 data datac

    table_number = num2str(f, '%01d');
    str = [directory_name, filename, table_number, '.csv'];
    path = join(str);

    data1 = readtable(path);

    data01(:,1) = data1(:,8); % Frame Number
    %data01(:,2) = data1(:,5); % x position
    %data01(:,3) = data1(:,6); % y position
    data01(:,2) = data1(:,17); % Total intensity

    data = table2array(data01);

    for d = 1:1:sectionnumber
        f1 = (d-1)*fps;
        f2 = f1 + fps -1;
        datac = data;
        data_tc = 0;
        intensity_bin = 0;
        a0 = 0;
        a1 = 0;
        a2 = 0;
```

```
a3 = 0;
a4 = 0;
a5 = 0;
a6 = 0;
a7 = 0;
a8 = 0;
a9 = 0;
a10 = 0;
a11 = 0;

for i = f1:1:f2

    V1 = find(datac(:,1) == i, 1, 'first'); %Find first particle in a frame
    V2 = find(datac(:,1) == i, 1, 'last'); %Find last particle in a frame

    checkempty = data_tc==0;

    if isempty (V1) == 0
        if checkempty == 1
            data_tc = datac(V1:V2,:);
        else
            m = size(data_tc,1);
            data_tc(m+1:m+V2-V1+1,:) = datac(V1:V2,:);
        end
    end

end

checkempty_1 = data_tc == 0;

if checkempty_1 ~= 1
    intensity_bin = data_tc(:,2);
    intensity_bin = intensity_bin*factor;

    for h = 1:1:size(intensity_bin)
        ins = intensity_bin(h);

        if ins < 860                                              % <26 nm
            a0 = a0+1;
        elseif ins > 860 && ins <= 2.0979e+03                     % 26~35 nm
            a1 = a1+1;
        elseif ins > 2.0979e+03 && ins <= 4.4588e+03              % 35~45 nm
            a2 = a2+1;
        elseif ins > 4.4588e+03 && ins <= 6.1163e+03              % 45~50 nm
```

```
                a3 = a3+1;
            elseif ins > 6.1163e+03 && ins <= 8.1408e+03      % 50~55 nm
                a4 = a4+1;
            elseif ins > 8.1408e+03 && ins <= 1.0569e+04      % 55~60 nm
                a5 = a5+1;
            elseif ins > 1.0569e+04 && ins <= 1.3438e+04      % 60~65 nm
                a6 = a6+1;
            elseif ins > 1.3438e+04 && ins <= 1.6783e+04      % 65~70 nm
                a7 = a7+1;
            elseif ins > 1.6783e+04 && ins <= 2.0642e+04      % 70~75 nm
                a8 = a8+1;
            elseif ins > 2.0642e+04 && ins <= 4.3742e+04      % 75~85 nm
                a9 = a9+1;
            elseif ins > 4.3742e+04 && ins <= 8.5256e+04      % 85~95 nm
                a10 = a10+1;
            else                                                      % >95
nm
                a11 = a11+1;
            end
        end

    end


    counts(d+sectionnumber*(f-1),1) = a0;
    counts(d+sectionnumber*(f-1),2) = a1;
    counts(d+sectionnumber*(f-1),3) = a2;
    counts(d+sectionnumber*(f-1),4) = a3;
    counts(d+sectionnumber*(f-1),5) = a4;
    counts(d+sectionnumber*(f-1),6) = a5;
    counts(d+sectionnumber*(f-1),7) = a6;
    counts(d+sectionnumber*(f-1),8) = a7;
    counts(d+sectionnumber*(f-1),9) = a8;
    counts(d+sectionnumber*(f-1),10) = a9;
    counts(d+sectionnumber*(f-1),11) = a10;
    counts(d+sectionnumber*(f-1),12) = a11;



  end


end
```

```
for col = 1:1:12

    counts_temp = counts(:,col);

    for row = 1:1:sectionnumber

        if row == 1
            counts_sum(row) = counts_temp(1);
        else
            counts_sum(row) = counts_sum(row-1) + counts_temp(row);
        end

        counts_evolve(row,col) = counts_sum(row);

    end

end
```

**Selective TRPV1 ion channel activation by local heat on W-PTM**

The Poly-D-lysine (PDL) (Sigma-Aldrich, Product Number: P6407) modified Au chip was obtained by culturing PDL solution (0.1 mg/mL) on Au chip in 37 °C for 2 h. After washing excessive PDL with PBS, the TRPV1 transfected HEK-293T cells were passaged on the Au chip and attached and grow overnight. Then the Fluo-4AM staining was performed. Meanwhile, extracellular solution (NaCl: 8.182 g; KCl: 0.402 g; $CaCl_2$: 0.294 g; $MgSO_4$: 1.204 g; HEPES: 1.191 g; Glucose: 1.802 g, water: 1 L) was prepared for keeping the osmotic pressure and healthy status of the cells during recording. After staining, a simple flow channel was quickly built on the gold chip with cells using double-sided tape and cover glass on the top. Then the extracellular solution was flowed on the chip at a steady speed of 200 µL/min controlled by a pressure pump.

P-polarized 660 nm laser at different power densities (0.50 and 1.2 $kW/cm^2$) were used to active the TRPV1 channels of the cells, while the fluorescence change at 515 nm was monitored by a second camera, excited by a 488 nm LED light.

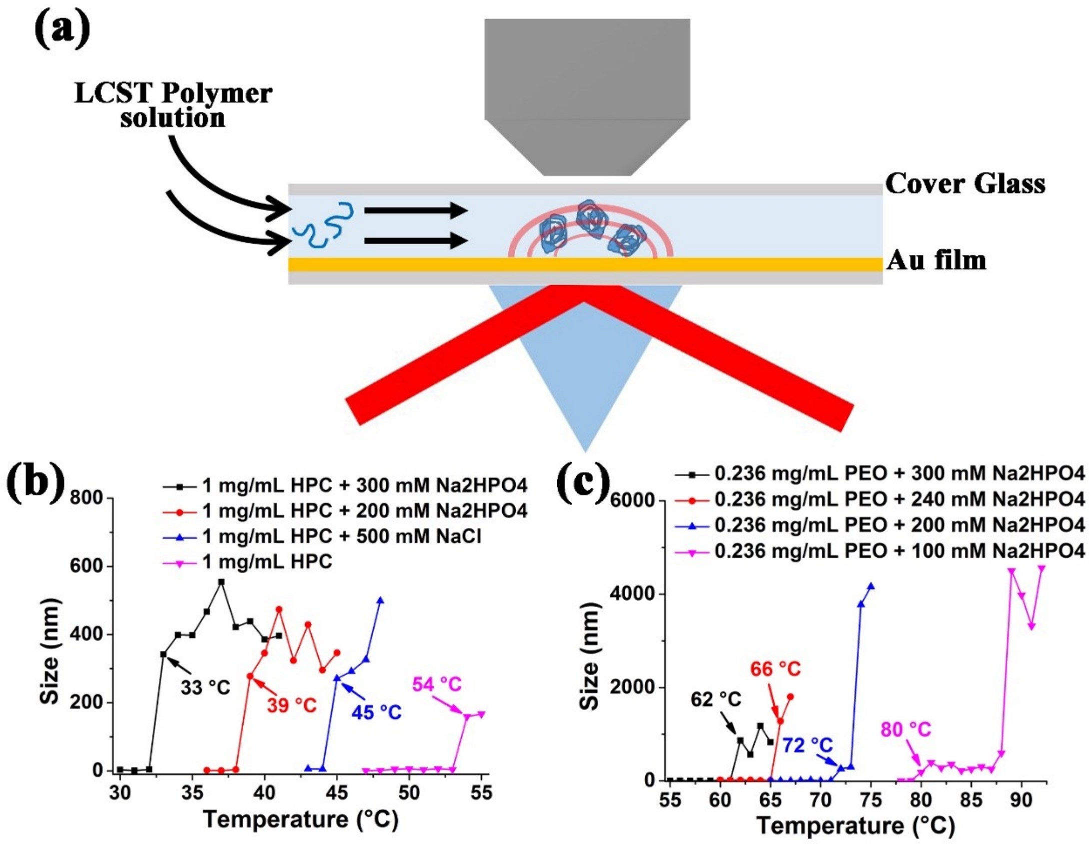


Figure S1. (a) Experimental set-up for the local temperature calibration of W-PTM using LCST polymer. (b, c) Phase transition temperature of the LCST polymers measured by DLS.

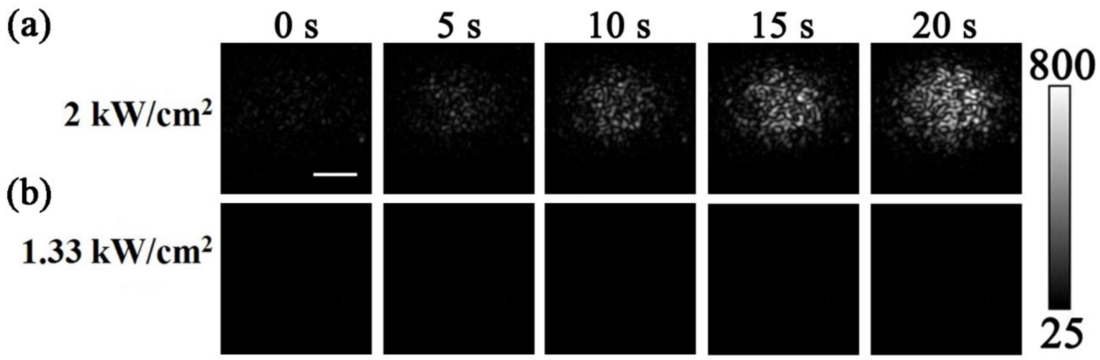


Figure S2. W-PTM imaging snapshots of 1 mg/mL HPC (LCST = 54 °C) excited at 2 kW/cm$^2$ (a) and 1.33 kW/cm$^2$ (b), respectively.

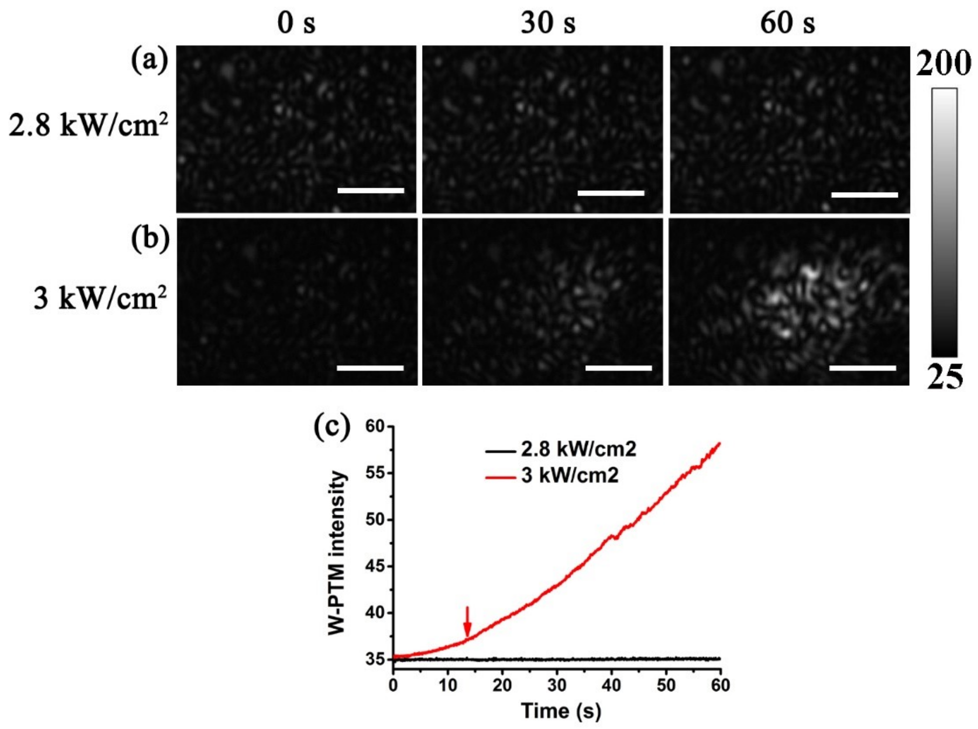


Figure S3. W-PTM imaging snapshots of 0.236 mg/mL PEO in 100 mM $Na_2HPO_4$ (LCST = 80 °C) excited at 2.8 kW/cm$^2$ (a) and 3 kW/cm$^2$ (b), respectively. (c) The ensemble W-PTM intensity as a function of excitation time in (a) and (b). The background fluctuation is around 0.3, so the phase transition starting point shall be defined when the W-PTM intensity is 1.5 higher than the starting value (indicated by the red arrow in (c)).

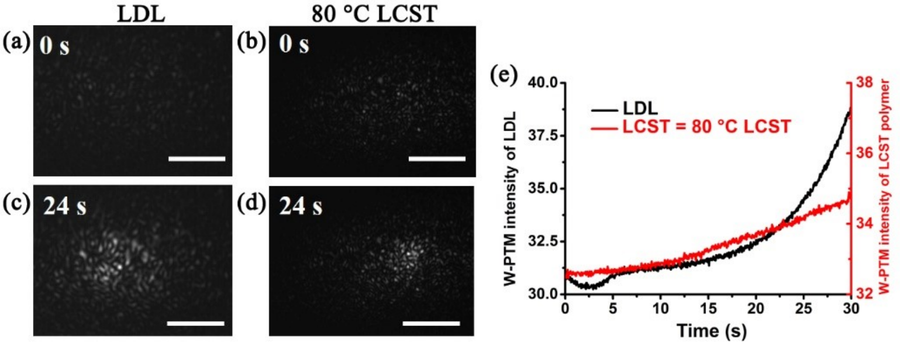


Figure S4. W-PTM imaging snapshots of 50 nM LDL aqueous solution (a, c) and 0.236 mg/mL PEO in 100 mM $Na_2HPO_4$ (80 °C LCST) (b, d), respectively. The excitation power density was 3 kW/cm$^2$ in both cases. Scale bar, 5 μm. (e) The ensemble W-PTM intensity as a function of excitation time for LDL and 80 °C LCST.

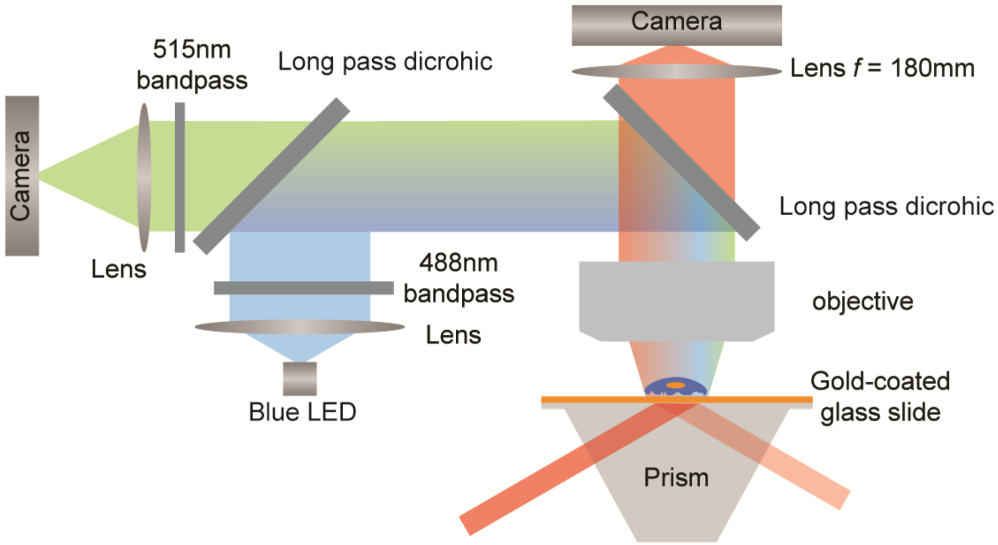


Figure S5. A detailed imaging set-up for selective TRPV1 activation monitoring by coupling FL imaging with W-PTM.

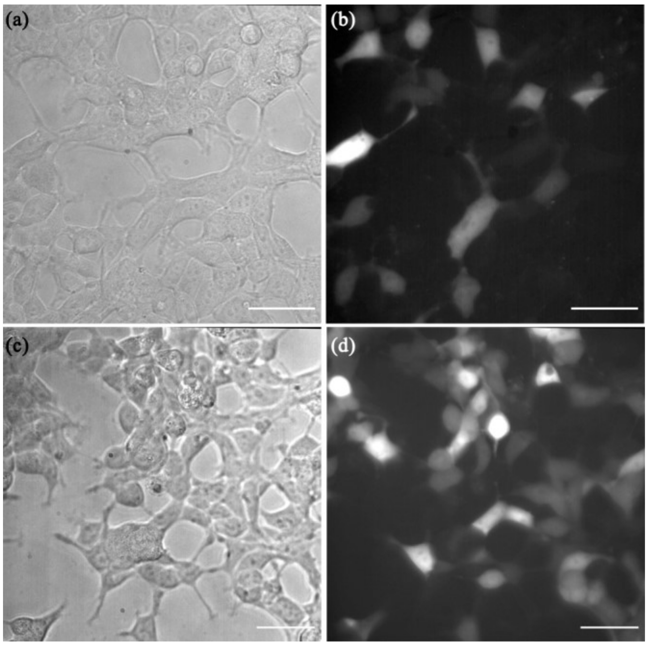


Figure S6. (a, b) HEK-293T cells transfected with TRPV1 for 24 h; (c, d) these cells were then transfected a second time in the same condition. (a, c) show the bright-field images; (b, d) are the corresponding fluorescence images excited at 525 nm. Acquisition time = 200 ms. The fluorescence of these cells comes from the dTomato indicator incorporated in the plasmid, which was transfected together with TRPV1. Thus its fluorescence indicates the TRPV1 transfection efficiency. It can be seen the transfection efficiency was improved from ~40% to ~90% after repeated transfection. Scare bar = 20 μm.

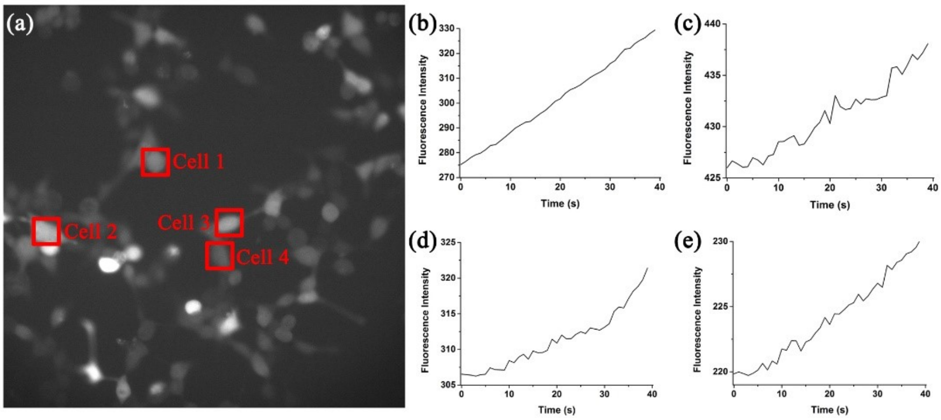


Figure S7. Fluorescence imaging monitored TRPV1 activation by adding capsaicin. (a) A snapshot of Fluo-4AM stained TRPV1 cells after adding capsaicin aqueous solution to the culturing media with a final concentration of 30 μM. Four cells of interest are marked in (a). (b-e) are corresponding fluorescence curves against treating time of these four cells. Excitation wavelength is 488 nm, acquisition time is 50 ms.
